# Does prefrontal connectivity during task switching help or hinder children’s performance?

**DOI:** 10.1101/2022.10.04.510761

**Authors:** Sina A. Schwarze, Corinna Laube, Neda Khosravani, Ulman Lindenberger, Silvia A. Bunge, Yana Fandakova

## Abstract

The ability to flexibly switch between tasks is key for goal-directed behavior and continues to improve across childhood. Children’s task switching difficulties are thought to reflect less efficient engagement of sustained and transient control processes, resulting in lower performance on blocks that intermix tasks (sustained demand) and trials that require a task switch (transient demand). Sustained and transient control processes are associated with frontoparietal regions, which develop throughout childhood and may contribute to task switching development. We examined age differences in the modulation of frontoparietal regions by sustained and transient control demands in children (8–11 years) and adults. Children showed greater performance costs than adults, especially under sustained demand, along with less upregulation of sustained and transient control activation in frontoparietal regions. Compared to adults, children showed increased connectivity between the inferior frontal junction (IFJ) and lateral prefrontal cortex (lPFC) from single to mixed blocks. For children whose sustained activation was less adult-like, increased IFJ-lPFC connectivity was associated with better performance. Children with more adult-like sustained activation showed the inverse effect. These results suggest that individual differences in task switching in later childhood at least partly depend on the recruitment of frontoparietal regions in an adult-like manner.

## 1. Introduction

The ability to switch flexibly between tasks enables individuals to adapt their behavior to changing environments and is a key component of cognitive control (Miyake et al., 2000; Miyake and Friedman, 2012). However, this flexibility comes at a cost. When faced with the demand to switch rapidly between tasks, individuals exhibit two types of performance decline. First, they show greater error rates and longer response times (RTs) on blocks of intermixed tasks (*mixed blocks*) compared to *single blocks* in which they perform only one task. These *mixing costs* are assumed to reflect sustained control demands, including the selection of the relevant goal (Chevalier et al., 2018; Chevalier and Blaye, 2009; Emerson and Miyake, 2003), and the maintenance and monitoring of multiple sets of rules associated with each task – so-called task sets (Braver et al., 2003; Pettigrew and Martin, 2016; Rubin and Meiran, 2005). Within the mixed blocks, performance declines further on trials that require a different task than the previous trial (i.e., *switch trials*) compared to trials repeating the previous task (i.e., *repeat trials*). These *switch costs* are thought to reflect transient control processes of inhibiting the no-longer-relevant task set (Allport et al., 1994; Meiran, 1996; Wylie and Allport, 2000) and retrieving and updating the newly relevant task set (Mayr and Kliegl, 2000; Rogers and Monsell, 1995).

Functional magnetic resonance imaging (fMRI) studies have highlighted the left inferior frontal junction (IFJ) and the left superior parietal lobe (SPL) as key regions involved in task switching (Derrfuss et al., 2005; Kim et al., 2012; Niendam et al., 2012; Richter and Yeung, 2014; Worringer et al., 2019; Zhang et al., 2021), particularly for transient processes including task-set updating and attentional shifts (Brass et al., 2005; Brass and von Cramon, 2004; Braver et al., 2003; Gurd et al., 2002). Sustained control processes, including task-set maintenance and monitoring, have been associated with the anterior and dorsolateral prefrontal cortex (aPFC and dlPFC, respectively; Badre, 2008; Braver et al., 2003; Gold et al., 2010) and the SPL (Brass and von Cramon, 2004; Bunge et al., 2003). Functional connectivity analyses have associated a frontoparietal network (FPN) of similar regions (i.e., the IFJ, SPL, dlPFC, and the precuneus) with transient control, and a cingulo-opercular network including the anterior insula/frontal operculum (aI/fO), the dorsal anterior cingulate cortex (dACC), and the aPFC with sustained control (Dosenbach et al., 2008, 2007, 2006).

Flexibly switching between tasks entails larger costs in children than adults (Cragg and Chevalier, 2012; Gupta et al., 2009; Huizinga et al., 2006; Huizinga and van der Molen, 2007). While children show similar switch costs to adults by 9-11 years, mixing costs only approach adult levels during adolescence (13–15 years; Cepeda et al., 2001; Crone et al., 2004, 2006a; Reimers and Maylor, 2005). The age differences reported previously suggest that sustained and transient control follow different developmental paths, with a more protracted development for sustained control, presumably reflecting the later maturation of relevant frontal brain regions and their connectivity.

The majority of developmental neuroimaging studies of task switching support a view of quantitative age differences, with children recruiting the same set of regions as adults, albeit in a less specific manner (Bunge and Wright, 2007; Engelhardt et al., 2019; Velanova et al., 2008; Wendelken et al., 2012; Zhang et al., 2021; but see Crone et al., 2006b; Morton et al., 2009). According to this view, the neural underpinnings of task switching typically observed in adulthood are in place by middle childhood and are subsequently fine-tuned, showing more differential activation between task conditions (Durston et al., 2006; Luna et al., 2015; Satterthwaite et al., 2013).

Similarly, by the age of around 10 years, children show network organizations that are similar to those of adults, but with continuing changes in connectivity strengths within and between networks (Engelhardt et al., 2019; Grayson and Fair, 2017; Marek et al., 2015). To date, only one study has examined age differences in network connectivity during task switching (Ezekiel et al., 2013). Children showed lower connectivity within the FPN than adults along with greater integration of the aPFC into the FPN, as reflected in ICA-based voxel-wise factor loadings, suggesting that children might configure brain networks in different ways than adults to meet increased control demands during task switching. However, this study did not examine age differences in connectivity change with increased control demand. Thus, it is unclear whether developmental changes in connectivity in response to increased control demands consist in the gradual evolution of the adult pattern, or whether one can observe a shift from a child pattern to an adult pattern. Conceptually, the brain typically provides more than one pathway to implement a task, both within and across individuals (Edelman, 1987; Lautrey, 2003; Li and Lindenberger, 2002). In the course of brain maturation, one way of implementing cognitive control may gradually become more effective than another, such that children eventually shift from one configuration to another (e.g., Van der Maas and Molenaar, 1992).

Of note, children differ considerably in task-switching development (Dauvier et al., 2012; Fields et al., 2021) and the pace of brain maturation (e.g., Mills et al., 2021). Church et al. (2017) reported that children who showed more adult-like (i.e., greater) frontoparietal activation during the preparatory period of task switching showed less activation in other regions during the target stimulus. Thus, age changes in brain-behavior mappings during task switching may not be uniform but follow different routes in different groups of children (e.g., Lautrey, 2003), creating an additional source of heterogeneity in the neural implementation of task-switching demands among children.

We examined the neural correlates of sustained and transient control during task switching in children aged 8 to 11 years and adults aged 20 to 30 years. We expected greater mixing and switch costs in children, reflecting the continued development of task-switching. We predicted that children would recruit similar frontal and parietal regions as adults but that they would show less upregulation of activation and connectivity in these regions with increased control demands. Further, we predicted that age differences in behavior and task-related activation would be more pronounced for sustained than transient control, consistent with the proposition that children reach mature levels of sustained control, operationally defined as task-set maintenance and monitoring, at a later age than mature levels of transient control, defined as task-set retrieval and updating. Finally, we explored whether individual differences in sustained activation and connectivity are associated with individual differences in task-switching performance. In particular, we were interested in the degree to which the link between connectivity and task-switching performance was modulated by the maturational status of individual children. We hypothesized that children with less mature sustained activation in the core task-switching network might show heightened connectivity with other brain regions, as a vicarious neural implementation of task-switching behavior.

## 2. Methods

### 2.1 Research Participants

Children between 8 and 11 years of age (n = 117; mean age = 10.0, SD = 0.71; 59 girls) and adults between 20 and 30 years of age (n = 53; mean age = 24.7, SD = 2.6; 28 women) were recruited from the internal participant database of the Max Planck Institute for Human Development. Participants were screened for MRI suitability, had no history of psychological or neurological diseases, and German was their primary language. All participants were right-handed. Adult participants and the participating children’s parents provided informed consent; the participating children additionally provided written assent. All participants were reimbursed with 10 € per hour spent at the MRI laboratory. The study was approved by the ethics committee of the Freie Universität Berlin and conducted in line with the Declaration of Helsinki.

Based on previous research on the developmental trajectories of sustained processes (measured by mixing costs) and transient processes (measured by switch costs) during task-switching (e.g., Cepeda et al., 2001; Crone et al., 2006a, 2004; Reimers and Maylor, 2005), we were particularly interested in the neural mechanisms supporting task switching between 8 and 11 years. This age group represents a period in childhood when both sustained and transient processes are still developing but are starting to dissociate, with transient control beginning to plateau and sustained control continuing to improve.

Participants were included in analyses if they performed the task in accordance with the task rules and had good fMRI data quality. Accuracy below 50% in the run of single blocks (run 1, see below for more details on the paradigm) or accuracy below 35% in either of the two runs of mixed blocks (run 2 and 3) was defined as poor performance. We defined a threshold well above chance (33%) in the single runs to make sure participants knew and could successfully apply the rules when no additional demand on switching among them was present. fMRI volumes with framewise displacement (Power et al., 2012) above 0.4 mm were marked as low-quality (see Dosenbach et al., 2017). If any of the fMRI runs exceeded 50% of low-quality volumes, the participant was excluded from further analysis. Twenty children were excluded because of excessive motion, four children due to poor performance, and four children due to both excessive motion and poor performance. Thus, a total of 89 children (mean age = 10.06, SD = 0.7, 50 girls) and 53 adults were retained in the analyses reported below.

### 2.2 Experimental Design

Participants performed a task-switching paradigm consisting of three different categorization rules (Figure 1A). In every trial, a face, a scene, and an object were presented, and participants were cued by the shape of the background (diamond, circle, or hexagon) as to whether to perform the face task, scene task, or object task. If the stimuli were presented in a diamond-shaped background, the face image had to be classified according to the age of the face (child vs. young adult vs. older adult). If the stimuli were presented in a circle, the image of the scene had to be classified according to the depicted scenery (forest vs. desert vs. sea). For stimuli presented in a hexagon, the image of the object had to be classified according to its color (yellow vs. red vs. purple). All stimuli and the task cue appeared at the same time. The arrangement of images on the screen varied randomly independent of the categorization rule. For each of the three tasks, participants had to respond via button press using their right index, middle, and ring finger. The assignment of finger to response option was held constant across participants. There were two different stimuli for each of the three levels of each of the three categorization rules, e.g., two faces of children, two yellow bags, two forest scenes, etc., resulting in a total of 18 different stimuli.

**Figure 1.**
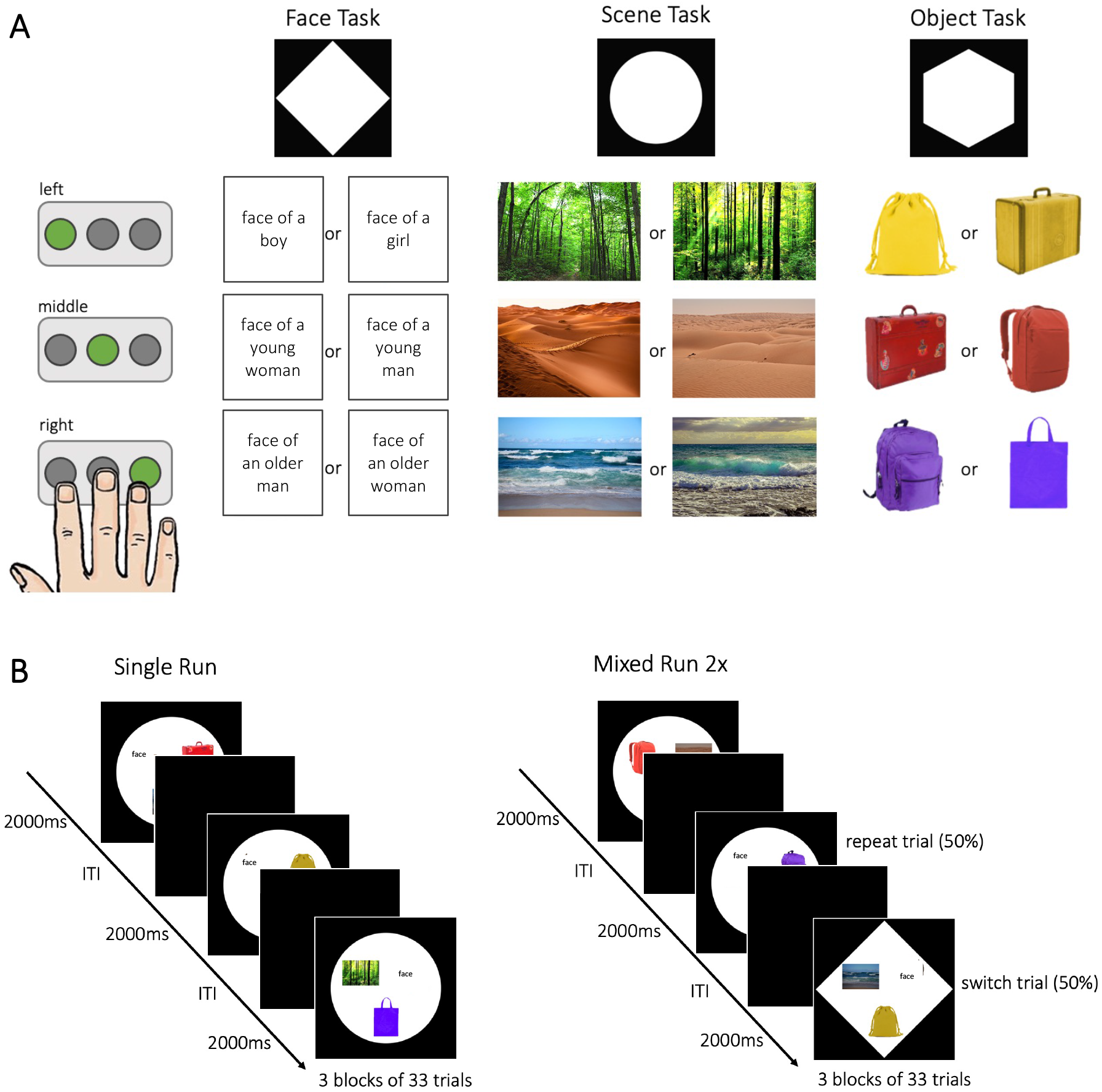
Experimental Design. (A) The task-switching paradigm consisted of three tasks, the face task, the scene task, and the object task. Participants had to perform the task indicated by the shape of the background. Depending on the stimulus presented, one of three buttons had to be pressed in response (here indicated by the green button). (B) Participants performed one single run, where the three tasks were presented in blocks of 33 trials and two mixed runs, where the tasks were intermixed. Each run consisted of 99 trials presented for 2 s with a jittered inter-trial interval (ITI). For a full description of the task, see text.

Participants performed three runs of 99 trials each (Figure 1B). Each trial lasted 2000 ms, followed by a fixation cross for a jittered time period (1000–6000 ms). After every 33 trials, there was an extended fixation period (20 s), resulting in three blocks of 33 trials per run. Each of the three runs lasted ca. 7 minutes. In the first run (i.e., *single blocks*), each block contained trials of a single task (i.e., the first 33 trials were of the face task, followed by 33 trials of the scene task, and 33 trials of the object task). Across runs two and three (i.e., *mixed blocks*), the three tasks were intermixed, so that participants needed to repeat a task in 50% of the trials (*repeat trial*) and switch to a different task in the other 50% of the trials (*switch trial*). The switches between task rules were not predictable, such that participants did not know in advance which task had to be performed next. In run two, 50 trials required a task switch and 48 a task repetition; in run three, 48 trials were switch trials and 50 were repeat trials. As the first trial of each run could not be categorized as a switch or repeat trial, it was excluded from all analyses.

The data analyzed here are part of a larger training study, during which participants were scanned multiple times. The present data were acquired during the second session of the training study. The first session of the study included a computerized assessment of general cognitive functioning not analyzed here, as well as a shortened version of the MRI task to ensure that participants were familiar with the rules. Prior to scanning, participants were shown the instructions again and practiced the task in a mock MRI scanner. The mock scanner looked identical to a regular MRI scanner, with MR scanning sounds being mimicked via speakers so that participants could get accustomed to the scanning environment. The three runs of the MRI task were performed in the scanner after an initial T1-weighted scan during which participants watched a muted cartoon.

### 2.3 Behavioral Analysis

Trials with response times (RT) below 200 ms and above 3000 ms were excluded from all analyses. Median RTs and mean accuracies were analyzed for effects of task rule condition (single block trials vs. repeat trials in mixed blocks vs. switch trials in mixed blocks), effects of age group (adults vs. children), and their interaction using separate linear mixed-effects models with a random intercept for subject. Only correct trials were considered for the RT analysis. Accuracy was calculated as the percentage of correct responses across all responses for a given condition. Outliers were defined as mean accuracy or median RTs that deviated by 3.5 standard deviations or more from the age-group mean, and were removed from analyses of accuracy and RT separately (Tabachnick and Fidell, 2013).

Main effects of condition were followed up by pairwise comparisons, with p-values FDR-corrected for multiple comparisons (Benjamini and Hochberg, 1995). Analyses were performed in R 4.0.3 (R Core Team, 2018) with the tidyverse (Wickham et al., 2019), ggpubr (Kassambara, 2020a), rstatix (Kassambara, 2020b), rmisc (Hope, 2013), sjPlot (Lüdecke et al., 2021), lme4 (Bates et al., 2022), lmerTest (Kuznetsova et al., 2017), and emmeans (Lenth et al., 2022) packages.

### 2.4 fMRI Data Acquisition and Preprocessing

Anatomical and functional MR images were collected on a 3-Tesla Siemens Tim Trio MRI scanner with a 32-channel head coil at the Max Planck Institute for Human Development. A high-resolution T1-weighted structural image was acquired (220 slices; 1 mm isotropic voxels; TE = 2.35 ms; FoV = 160 x 198 x 220). Functional runs consisted of 230 whole-brain echo-planar images of 36 interleaved slices (TR = 2000 ms; TE = 30 ms; flip angle = 80°; slice thickness = 3 mm; in-plane resolution (matrix) = 3 x 3 mm (72 x 72); FoV = 216 x 216 x 129.6). During the acquisition of functional images, the task was projected on a screen behind the participant’s head that they could see via a mirror mounted on the head coil. During task execution participant motion was monitored using Framewise Integrated Real-time MRI Monitoring (FIRMM; Dosenbach et al., 2017). Participants were provided feedback on their movement after each run and were encouraged to try to move less if they exceeded the threshold of 0.4mm of movement for at least 10% of frames of the previous run.

Preprocessing was performed using fMRIprep (Version 20.2.0; Esteban et al., 2019). For a detailed description, see the fMRIprep documentation (link). Briefly, BOLD images were co-registered to individual anatomical templates using FreeSurfer, which implements boundary-based registration (Greve and Fischl, 2009). Additionally, they were slice-time corrected (using AFNI; Cox and Hyde, 1997), and realigned (using FSL 5.0.9; Jenkinson et al., 2002). Finally, images were resampled into MNI152NLin6Asym standard space with a voxel size of 2 mm x 2 mm x 2 mm and spatially smoothed with an 8 mm FWHM isotropic Gaussian kernel using SPM12 (Functional Imaging Laboratory, UCL, UK).

### 2.5 fMRI Data Analysis

#### 2.5.1 General Linear Model

GLM analyses were performed using SPM12 software. For each subject, we estimated two general linear models (GLM). The first GLM modeled sustained control demands in a blocked design (*sustained* GLM). Condition regressors included *single task*, consisting of three blocks of 127 s each within the first run, and *mixed task,* consisting of six blocks of 127 s each, evenly split over the second and third run. Data were high-pass filtered at 172 s.

A second GLM modeled transient control demands in an event-related design (*transient* GLM). Here, each stimulus presentation was coded as an event with zero duration, and convolved with a canonical hemodynamic response function (HRF). Separate regressors were included for correct *switch trials* and *repeat trials*. Incorrect trials, trials with extreme RTs (below 200 ms or above 3000 ms), trials with missed responses, and the first trial of each run were modeled as nuisance regressors. Data were high-pass filtered at 128 s.

To minimize head motion artifacts, both GLMs included the amount of frame displacement per volume, in mm (Power et al., 2012), realignment parameters (three translation and three rotation parameters), and the first six anatomical CompCor components (as provided by fMRIprep; Behzadi et al., 2007) as regressors of no interest. The first five volumes of each run were discarded to allow for stabilization of the magnetic field and temporal autocorrelations were estimated using first-order autoregression.

Sustained control activation was defined as higher activation in mixed compared to single blocks (mixed > single) in the sustained GLM. Transient control activation was defined as higher activation on switch trials compared to repeat trials (switch > repeat) within the mixed runs of the transient GLM. The sustained GLM modeled control in a continuous manner, such that mixed blocks included both switch and repeat trials (e.g., Braver et al., 2003). With these separate models, it is not possible to directly compare single and repeat trials; thus, we also constructed an event-related GLM that included correct single, repeat, and switch trials as separate regressors of interest (*control* GLM). All nuisance regressors and additional parameters were identical to the transient GLM.

#### 2.5.2 Region-of-Interest Definition and Analysis

First, we identified regions of interest (ROIs) involved in sustained and transient control over all subjects (whole-brain analyses; p < .05 family-wise-error (FWE) corrected). For sustained control, we selected voxels that showed greater activation in mixed blocks than in single blocks in the sustained GLM. We identified regions involved in transient control by selecting voxels showing greater activation on switch than on repeat trials in the transient GLM. Second, to identify ROIs that showed modulation by sustained and transient control demand, which we referred to as *overlap* ROIs, we applied inclusive masking which identified common voxels for sustained and transient activation. Finally, to define regions that showed modulation by either sustained or transient control demand, we applied the map of the overlap as an exclusive mask to the sustained control contrast—which we termed *sustained* ROIs—and the transient control contrast, termed *transient* ROIs. Only clusters of at least 50 voxels were considered for ROI definition. If clusters were stretched across multiple anatomical areas, they were additionally masked using anatomical regions of the Harvard-Oxford atlas (Makris et al., 2006) thresholded at 30%. No anatomical mask is available for the IFJ; therefore this ROI was created by splitting the cluster based on a meta-analysis (Derrfuss et al., 2005). Hemisphere masks were applied to restrict clusters that extended beyond one hemisphere. Finally, ROIs were binarized and used as masks for beta-parameter extraction using MarsBar (Brett et al., 2002).

With these ROI analyses, we sought to test whether children and adults differed in the extent of upregulation of sustained and transient control activation with increased control demand. Upregulation of sustained control was computed as the difference between parameters of mixed and single blocks, and upregulation of transient control was computed as the difference between parameters of switch and repeat trials. We then compared the extent of upregulation between age groups for each ROI using t-tests in R, reporting Cohen’s d as effect size (Cohen, 1988) and p-values FDR-corrected for multiple comparisons.

To examine age differences in regions involved in task switching that were not defined in the present task, we conducted additional analyses in ROIs based on a metaanalysis in children and adults (Zhang et al., 2021). Specifically, we constructed 6 mm spheres around the activation peaks for task switching across both age groups (see Supplementary Table 2). Subsequently we extracted activation parameters from the *sustained* and *transient* GLMs and tested for age differences using t-tests as described above.

To investigate whether children or adults recruited additional brain regions beyond the ones we identified over both age groups, we computed exploratory whole-brain age comparisons of the mixed > single and switch > repeat contrasts in the sustained and transient GLM, respectively (p < .05, FWE-corrected).

#### 2.5.3 Psychophysiological Interaction Analysis

Generalized psychophysiological interactions (gPPI; McLaren et al., 2012) were analyzed using the CONN toolbox (Version 20b; Whitfield-Gabrieli and Nieto-Castanon, 2012). Unlike correlational measures of functional connectivity, PPI models how the strength of coupling between a seed region and a voxel elsewhere in the brain differs across conditions. To this end, a GLM predicting the time series of the target voxel by the interaction regressor is constructed. The interaction regressor is created by multiplying the time series of the seed region with the task regressor for each condition (i.e., the condition onset times convolved with the HRF). The PPI analysis focused on effects of sustained control, as we had predicted greater age differences here.

The main effects of the two sustained control conditions (single and mixed blocks) and the effect of the nuisance regressors previously applied to the sustained GLM were regressed from the BOLD signal time series before estimating the interaction factor. Importantly, voxel-level time series were estimated using the smoothed data, while ROI level (i.e., seed) time series were estimated using unsmoothed data to prevent a “spillage” of the BOLD signal of voxels outside the ROI into the ROI time series. Interaction parameters were estimated separately for each condition, i.e., single and mixed blocks (McLaren et al., 2012).

Based on the prominent role of the left IFJ in task switching (Derrfuss et al., 2005; Kim et al., 2012; Richter and Yeung, 2014) and the fact that it showed activation in our univariate analysis, we selected it as a seed region for the PPI analysis. More specifically, a sphere of 6 mm radius was placed around the group peak coordinates of the mixed > single contrast of the sustained GLM.

We compared the difference in seed-to-whole brain connectivity for the IFJ for mixed vs. single blocks between children and adults. Analyses were performed on a gray-matter mask of the whole brain and corrected for multiple comparisons at the cluster level (p < .05 FDR-corrected, voxel threshold at p < .001 uncorrected). Beta estimates were extracted from clusters showing age differences in connectivity for the mixed > single contrast to visualize the results and to relate connectivity patterns to performance in the task. Additionally, the large cluster in the prefrontal cortex (PFC) was split into a medial and lateral part, as these have been associated with different aspects during task switching (Koechlin et al., 2000). More specifically, the medial PFC (mPFC) was defined by the superior frontal and paracingulate gyrus and the lateral PFC (lPFC) by the frontal pole and middle frontal gyrus of the Harvard-Oxford Atlas (Makris et al., 2006), thresholded at a probability of 30%.

#### 2.5.4 Functional Activation Deviation Scores

To take individual differences among children in sustained activation into consideration, we computed functional activation deviation scores (cf. Düzel et al., 2011; Fandakova et al., 2015). The deviation scores reflect the similarity of task-related activation of an individual child to the adult mean. To this end, we created a mask including all clusters that showed sustained activation across adults (p < .05, FWE-corrected). The deviation score for each child was then calculated as the difference between the average T-value of the sustained control contrast (mixed > single) of all voxels inside this adult mask and outside it. Thus, a negative score represented more adult-like activation patterns, and a more positive score represented less adult-like patterns.

To examine relations between deviation scores, connectivity, and performance, we used the adjusted accuracy mixing costs. Adjusted mixing costs were calculated based on a measure of task-switching efficiency (Brüning and Manzey, 2018) by dividing the difference between single- and mixed-block accuracy by the single-block accuracy, and thus allowed us to adjust mixing costs to differences in overall performance. The adjustment to the mixing costs was applied because children varied considerably in their single-block performance, with about 12% of children showing single-task accuracy below 80%. We focused on individual differences in accuracy rather than RT, as we only observed age group differences in mixing and switch costs in accuracy. We posit that this study yielded lower accuracy because the task was more challenging than in prior studies, in that it required switches between three tasks with three stimulus-response mappings each, as opposed to, for example, two tasks with two arbitrary mappings each (e.g., Crone et al., 2006a; Reimers and Maylor, 2005). As a measure of connectivity, we used the difference in PPI beta parameters between mixed and single blocks of the connections between the IFJ and the lateral and medial PFC clusters separately, to investigate whether they showed different influences on performance.

## 3. Results

### 3.1 Greater Costs of Task Switching in Children

Accuracy results are shown in Figure 2A. We fit a linear mixed-effects model with accuracy as the dependent variable and the factors age group (children vs. adults) and condition (single vs. repeat vs. switch), the interaction between these two factors and a random intercept for subject using log-likelihood optimization. To enable comparisons of mixing costs (i.e., performance differences between repeat and single trials) and switch costs (i.e., performance differences between switch and repeat trials), the condition repeat was used as the reference condition in the model. The model revealed a significant effect of age group (β = −0.068, p < .001) and the switch condition (vs. repeat; β = −0.031, p = .01). The effect for the single condition (vs. repeat) was at trend level (β = −0.024; p = .06). Thus, children showed overall lower accuracy (M = 0.81, SD = 0.16) than adults (M = 0.96, SD = 0.04) and across age groups that accuracy was highest on single trials (M = 0.94, SD = 0.08), followed by repeat trials (M = 0.87, SD = 0.13), and then switch trials (M = 0.80, SD = 0.17). There were significant interactions of age group with both conditions (single: β = −0.089, p < .001; switch: β = −0.054, p < .001) indicating that children showed greater differences between the conditions than adults, especially with respect to the difference between single and repeat conditions. Taken together, while both age groups demonstrated reliable mixing and switch costs, the age differences for mixing costs (i.e., the interaction of age group and single condition) were more pronounced than the age differences for switch cost (i.e., the interaction of age group and switch condition).

**Figure 2:**
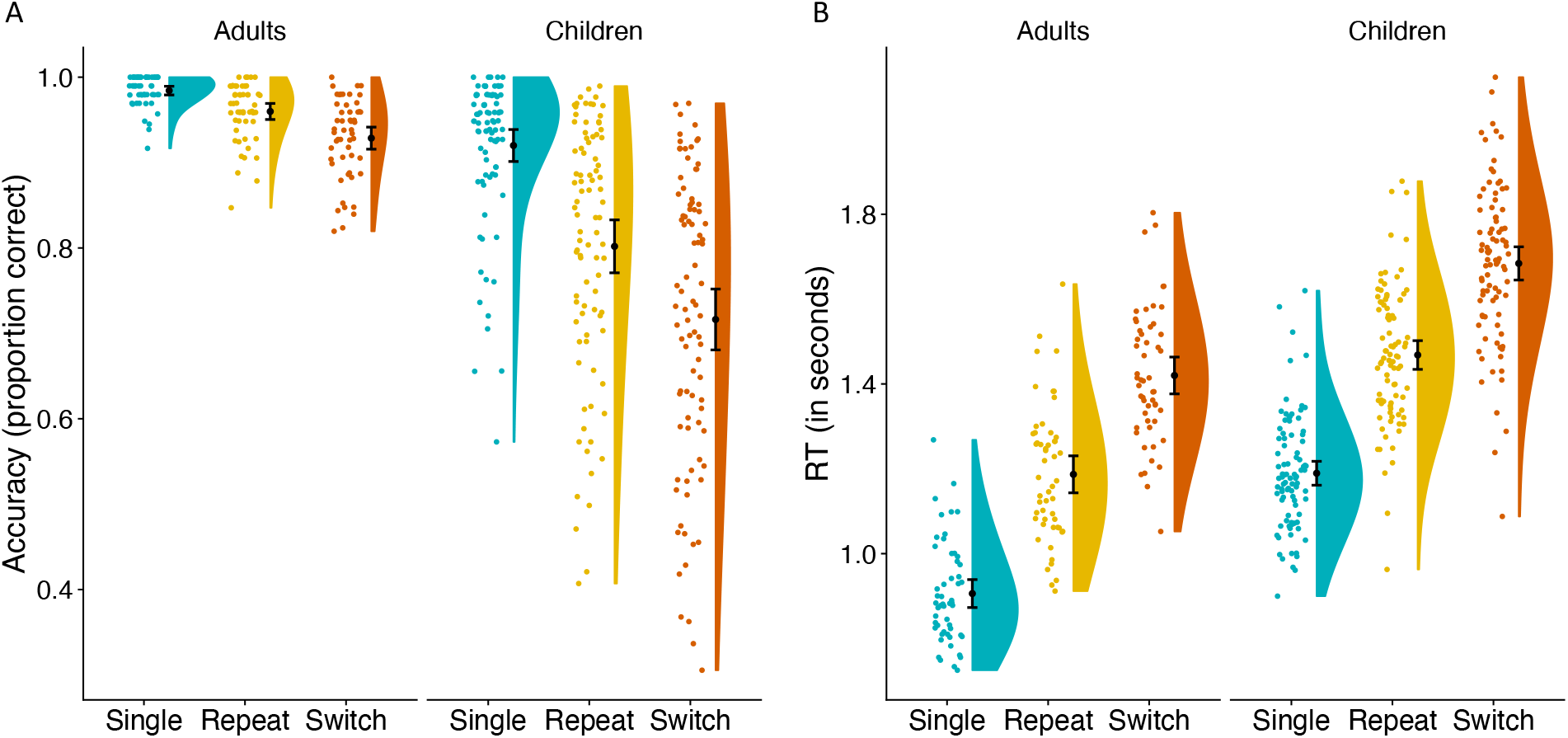
Behavioral results. (A) Proportion of correct responses for single, repeat, and switch trials. Across age groups accuracy was highest for single trials, followed by repeat and switch trials. Children showed greater mixing (single vs. mixed trials) and switch (repeat vs. switch trials) costs, while age differences were more pronounced for mixing costs. (B) Response time for single, repeat, and switch trials. Across age groups, responses to single trials were fastest, followed by repeat and switch trials. Mixing and switch costs in response times (RT) were similar across age groups.

RT results are shown in Figure 2B. The mixed-effects model of RT, set up with the same fixed and random effects as the model of accuracy, revealed significant effects of age group (β = 0.28, p < .001) and both conditions (single: β = −0.27, p < .001; switch: β = 0.24, p < .001; both compared to repeat as reference). There was no significant age-group-by-condition interaction, neither with the single condition (β = 0.0047, p = .83) nor with the switch condition (β = 0.013, p = .54). Overall, children (M = 1.45 s, SD = 0.26) responded more slowly than adults (M = 1.17 s, SD = 0.26). Across both groups, responses to single trials (M = 1.08 s, SD = 0.19) were fastest, followed by repeat (M = 1.35 s, SD = 0.20) and switch trials (M = 1.60 s, SD = 0.22).

Taken together, children and adults demonstrated reliable declines in performance both in terms of lower accuracy and slower RTs going from single blocks to mixed blocks, and from repeat trials to switch trials. Critically, children showed greater declines in accuracy than adults with increasing task switching demands, with age differences being particularly pronounced for mixing costs.

### 3.2 Frontal and Parietal Brain Regions Associated with Sustained and Transient Control

Analyses of whole-brain activation associated with sustained control (mixed > single blocks) across children and adults indicated enhanced activation in multiple bilateral prefrontal and parietal regions (see Supplementary Table 1), including the IFJ, SPL, dACC, and dlPFC (see Figure 3, green color). A whole-brain analysis of transient control (switch > repeat trials) across children and adults revealed enhanced activation in left lateral and medial parietal and prefrontal regions as well as bilateral occipital regions, including the left IFJ, left SPL, left dACC, and the bilateral precuneus (Figure 3, red color; Supplementary Table 1).

**Figure 3:**
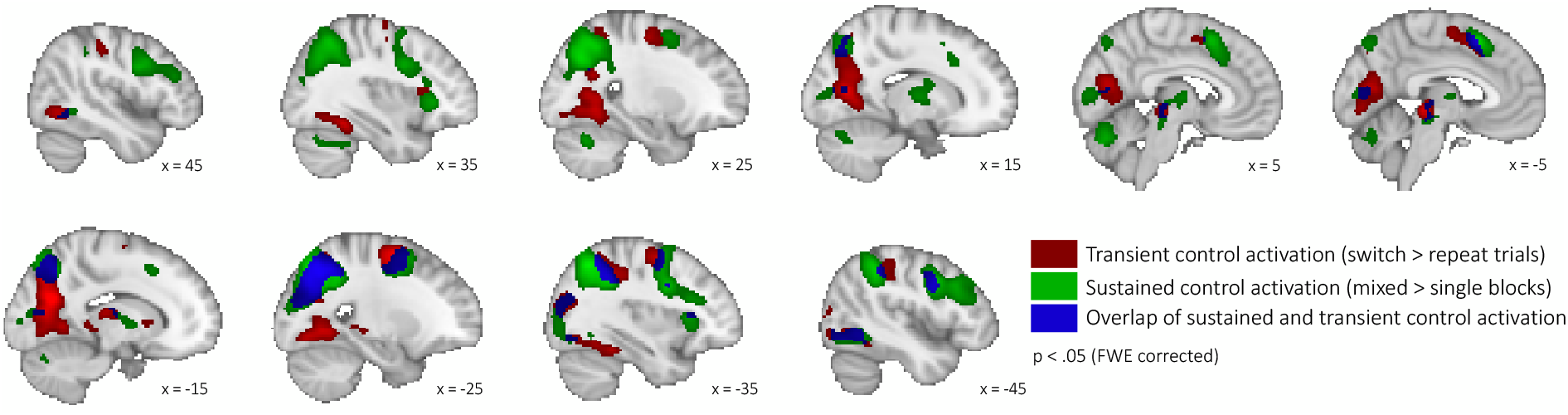
Univariate activation across age groups. Transient control activation (switch > repeat trials) depicted in red, sustained control activation (mixed > single blocks) in green, and overlap (voxels showing transient and sustained activation) in blue. N = 142 (53 adults, 89 children), p < .05 FWE-corrected, k = 50.

Activation patterns of sustained and transient control partially overlapped. Inclusive masking revealed several regions activated by both sustained and transient control demands, including the left IFJ, left SPL, left dACC, left inferior lateral occipital cortex (iLOC), left anterior insular (aI), and left middle frontal gyrus (MFG). On the other hand, regions exclusively showing modulation of activation with sustained control demand included the bilateral dlPFC, right IFJ, right SPL, right aI, and right MFG. Activation in the bilateral precuneus and a region in the bilateral premotor cortex (preMC) was exclusively modulated with transient control demand.

Thus, across age groups, we found frontal and parietal regions to be associated with sustained and/or transient control, including regions—the IFJ, SPL, dACC, dlPFC, and precuneus—that have been associated with task switching in previous research in adults (Kim et al., 2012; Richter and Yeung, 2014). Of these regions, the left IFJ, SPL, and dACC showed modulation of activation for both sustained and transient control demand, bilateral dlPFC exclusively for sustained control demand, and bilateral precuneus exclusively for transient control demand.

To test whether sustained control effects were driven by switch trials during mixed blocks or were also present when comparing single and repeat trials only, we performed a control GLM that included single, repeat, and switch trials in an event-related design. Univariate results of the repeat > single contrast of this control GLM were comparable to the mixed > single contrast of the sustained GLM (see Supplementary Figure 1), suggesting that sustained control effects were jointly driven by switch and repeat trials within the mixed blocks.

### 3.3 Less Upregulation of Sustained and Transient Control Activation in Children

Having identified regions across the whole sample that are involved in one or both types of control across both age groups, we next sought to examine age differences in the modulation of activation by sustained and transient control demands within these regions. We tested for age differences in the upregulation of sustained control activation by performing t-tests on the difference between mixed and single parameter estimates. As shown in Figure 4A, the overlap ROIs, including left IFJ, SPL, and dACC, showed a significant age difference in the degree of increase in activation from single to mixed blocks (IFJ: t(118.66) = −3.86, p_corrected_ < .001, Cohen’s d = 0.66; SPL: t(125.71) = −4.02, p_corrected_ < .001, d = 0.68; dACC: t(121.59) = −5.57, p_corrected_ < .001, d = 0.72; FDR-corrected), such that, across all ROIs examined, children showed less upregulation of activation from single to mixed blocks.

**Figure 4:**
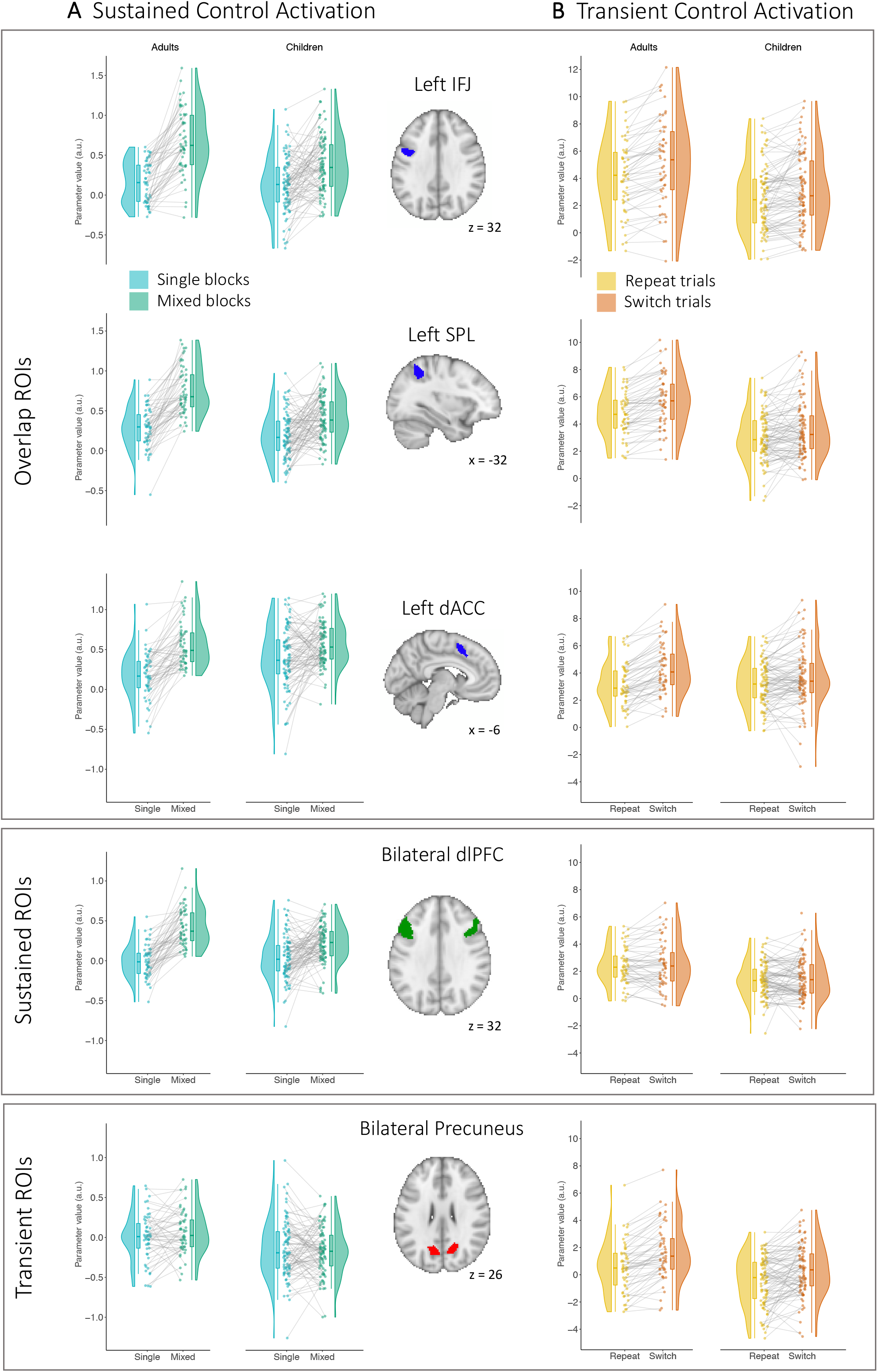
Age differences in activation due to sustained and transient control demands by ROI. (A) Extracted parameter estimates for single blocks (blue) and for mixed blocks (green). Children showed less upregulation of activation due to sustained control demand in the left IFJ, SPL, and dACC, and the bilateral dlPFC. Neither age group showed upregulation of sustained control activation in the Precuneus. (B) Extracted parameter estimates for repeat trials (yellow) and for switch trials (orange). Children showed less upregulation of activation due to transient control demands in the left IFJ, SPL, dACC, and the bilateral precuneus. Neither age group showed upregulation of transient control activation in the dlPFC.

Similarly, we found significant age differences in the extent of upregulation of activation by sustained control demands in the bilateral dlPFC, right IFJ, and right SPL, ROIs that were exclusively associated with sustained control (dlPFC: t(127.31) = −4.49, p_corrected_ < .001, d = 0.76; IFJ: t(107.1) = −3.12, p_corrected_ = .003, d = 0.54; SPL: t(122.58) = −2.96, p_corrected_ = .004, d = 0.5, FDR-corrected, see Figure 4A and Supplementary Figure 3), indicating that children showed less upregulation of task-related activation in the mixed blocks than adults. Taken together, these results suggest that even though children upregulated brain activity in regions that have been identified as key task-switching regions across both age groups with increasing demands on sustained control, they did so to a lesser extent than adults, both for overlap and sustained ROIs.

Next, we tested for age differences in the upregulation of transient control activation by performing t-tests on the difference between switch and repeat parameter estimates (see Figure 4B). The overlap ROIs in left IFJ, SPL, and dACC showed significant age differences in upregulation of transient control activation (IFJ: t(130.14) = −2.25, p_corrected_ = .045, d = 0.38; SPL: t(138.61) = −2.30, p_corrected_ = .045, d = 0.37, dACC: t(135.53) = −3.74, p_corrected_ = .002, d = 0.62, FDR-corrected). The bilateral precuneus, exclusively associated with transient control, also showed greater upregulation of transient control activation in adults than in children (t(134.18) = −2.32, p_corrected_ = .045, d = 0.39, FDR-corrected). Taken together, children upregulated activation due to transient control demand to a lesser extent than adults did across the key regions implicated in task-switching.

The preceding ROI-based analyses revealed age differences in activated regions common to adults and children. However, it is possible that children or adults showed different activation in additional regions. Thus, we also conducted an exploratory analysis investigating age differences in sustained and transient activation on the whole-brain level. Consistent with the results of the ROI analysis, the whole-brain analysis of sustained control showed greater upregulation of activation (mixed > single) for adults than children in the left SPL, left IFJ/dlPFC, and left superior lateral occipital cortex (p < .05 FWE-corrected; Supplementary Figure 4a). No regions showed greater upregulation of sustained control activation in children. Further, no regions showed age differences in upregulation of activation due to transient control demand (switch > repeat); for further details, see the Supplementary Material.

Finally, we conducted additional analyses in ROIs based on a recent meta-analysis of task switching across adolescents and adults (Zhang et al., 2021) in order to examine age differences in regions that were not specifically defined in the context of the present task. These additional analyses revealed a similar pattern as reported above. More specifically, we found significant age differences in activation associated with sustained control in multiple frontoparietal regions, including bilateral SPL, bilateral middle frontal gyrus, and precentral gyrus (see Supplementary Table 2 and Supplementary Figure 5). With regards to transient control, fewer regions showed age differences in activation, including two ROIs in the left and right middle frontal gyrus (see Supplementary Table 2 and Supplementary Figure 5). The remining regions (i.e., SPL, precentral gyrus, superior frontal gyrus) showed no significant differences between children and adults.

### 3.4 Greater Increases in Connectivity Under Increased Sustained Control Demand in Children

Based on previous research (e.g., Cepeda et al., 2001) showing that differences between adults and children in the age range investigated here are more pronounced during sustained control, and the fact that our results showed more pronounced age differences for mixing costs, we focused on connectivity associated with sustained control. More specifically, we were interested in the regions showing age differences in connectivity with the IFJ under increased sustained control demands. The left IFJ was selected as a seed, as it has been identified as a key brain region during task switching (Derrfuss et al., 2005; Kim et al., 2012; Richter and Yeung, 2014).

A whole-brain PPI analysis revealed that children showed a more pronounced increase in connectivity with the IFJ from single to mixed blocks in the bilateral angular gyrus (AG), and the medial and lateral prefrontal cortex (mPFC and lPFC, respectively) (p < .001, FDR cluster corrected p < .05). Neither of these regions overlapped substantially with the regions showing greater sustained control activation across age groups (see Supplementary Figure 6). The PFC cluster was split to account for possible differences between lateral and medial regions (Koechlin et al., 2000). The lPFC cluster was localized anteriorly adjacent to the bilateral dlPFC that showed sustained activation, and the mPFC cluster was localized anterior to the dACC showing transient and sustained activation. Figure 5 shows extracted connectivity parameters of the clusters that showed age differences in the comparison of mixed and single blocks. Specifically, age differences were driven by the fact that children showed higher IFJ–mPFC, IFJ–lPFC, and IFJ–AG connectivity in mixed than in single blocks, whereas adults did not show differences in connectivity for these regions between conditions. Additionally, Figure 5 illustrates the variation in the increase of connectivity from single to mixed blocks among children, such that some showed very steep increases, while others showed rather shallow increases or even decreases. Thus, these data suggest that many, but not all children showed a greater control demand-related increase in connectivity to broad swaths of prefrontal and parietal cortices outside the core areas implicated in task-switching. Note, however, that while these age group differences were identified in the aggregated group comparison, the individual estimates presented in Figure 5 suggested large interindividual differences, such that some children showed more pronounced increases in connectivity, while others showed smaller increases.

**Figure 5:**
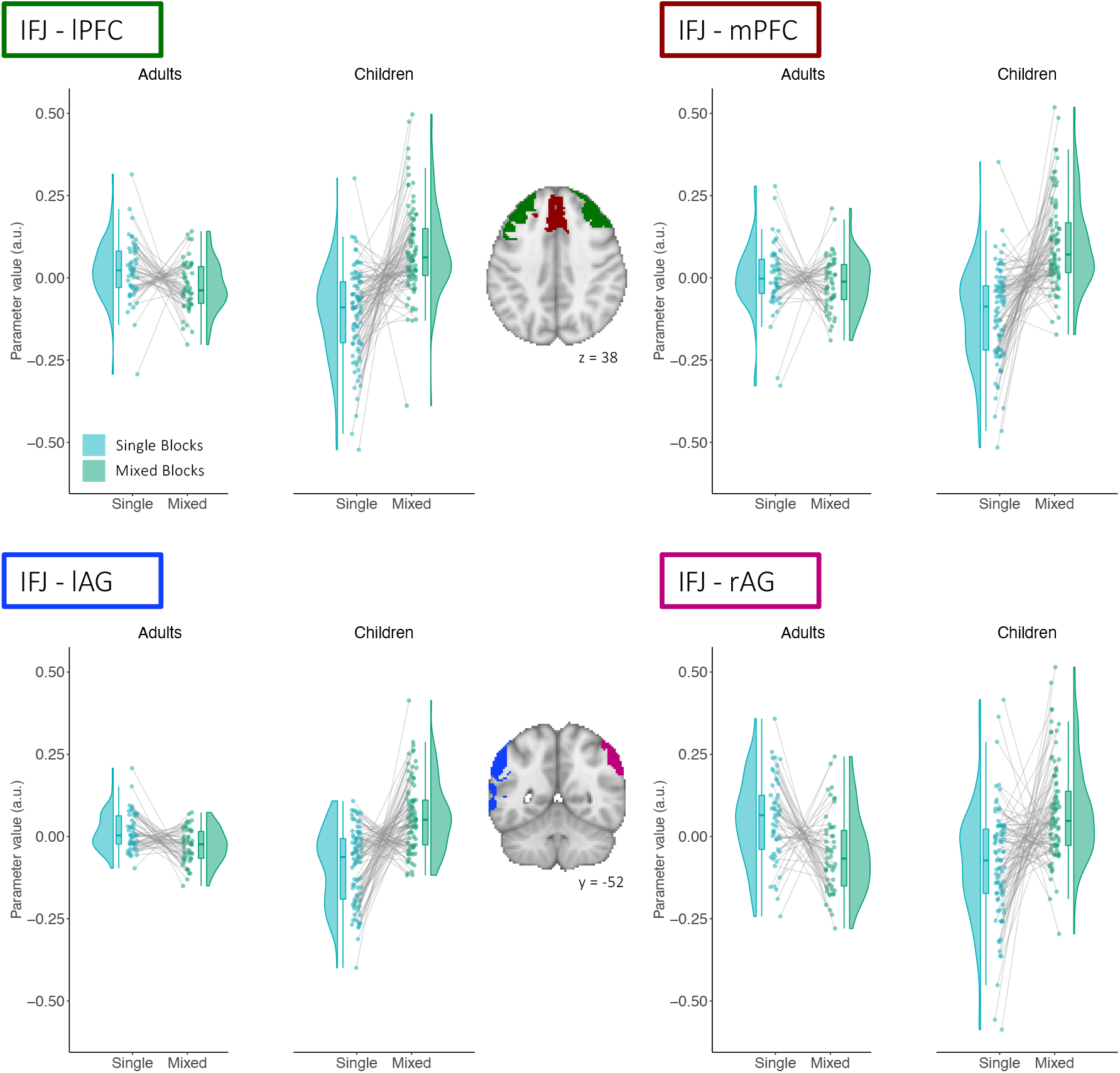
Age differences in connectivity increases under greater sustained control demands. PPI parameter estimates for clusters showing greater connectivity with the IFJ during mixed blocks (green) than during single blocks (blue) in children compared to adults identified in a whole-brain analysis with the left IFJ as a seed.

### 3.5 Connectivity Effects on Performance Depend on Adult-like Activation in Frontal and parietal Regions

Children showed less upregulation of activation in sustained control regions, but a greater increase in IFJ connectivity with additional regions as a function of increased demands on sustained control, including regions in lateral and medial PFC that have been implicated in cognitive control processes (e.g. Braver et al., 2003). This pattern of results raises the important question whether the broader connectivity pattern observed in children represents an alternative, presumably developmentally earlier way to support task performance specifically in those children whose frontoparietal activation patterns differ more markedly from those observed among adults. If so, the relation of connectivity increases and accuracy may vary across children, and may depend on the extent to which an individual child showed a mature pattern of sustained control activation.

To test this hypothesis, we computed deviation scores indicating how similar each child’s sustained control activation was to that of the average adult. For every individual child, this score was calculated by subtracting the average T-value of the mixed > single contrast of all voxels *inside* the clusters that showed sustained control activation in adults from the average T-value of the mixed > single contrast of all voxels *outside* of the clusters that showed such activation in adults (cf. Düzel et al., 2011; Fandakova et al., 2015). Thus, a more negative score indicates less deviation from the average adult activation pattern, while a more positive score indicates greater deviation and therefore a less adult-like activation pattern. This approach allowed us to take two previously observed aspects of age differences in brain activation into account. First, age differences have been found in the strength of task-related activation within the same regions, such that children show overall weaker activation than adults (e.g., Wendelken et al., 2012). Second, children have been shown to have less specific task-related activation than adults (e.g., Durston et al., 2006). Both of these aspects would result in more positive scores, indicating a greater difference to the average adult pattern. The deviation scores were not correlated with children’s age (r = −.088, p = .41). However, our sample was not evenly distributed across the age range (see Supplementary Figure 7), limiting our ability to adequately test for this association.

We performed a linear regression on the accuracy mixing costs with deviation scores, increases in IFJ–PFC connectivity (in response to increased sustained control demands), and the deviation score by connectivity interaction as predictors. The goal of these analyses was to examine whether the association between connectivity and performance depended on the maturity of activation under sustained control demands. Regressions were performed separately for connectivity with the lPFC and mPFC clusters.

We found significant main effects of deviation score (β = 0.07, p = .009) and of increased IFJ–lPFC connectivity (β = −0.18, p = .026) on mixing costs. However, these effects were qualified by a significant interaction between deviation score and connectivity (β = −0.22, p = .03). Thus, the association between connectivity increases and performance was modulated by the extent to which children showed an adult-like pattern of sustained control activation (Figure 6). Specifically, for children with a more adult-like activation pattern (i.e., more negative deviation scores), a greater increase in connectivity for mixed blocks was associated with greater mixing costs, or lower performance. Thus, for children with more adult-like activation patterns, the inclusion of the lPFC in the control network due to greater sustained control demands was associated with poorer performance.

**Figure 6:**
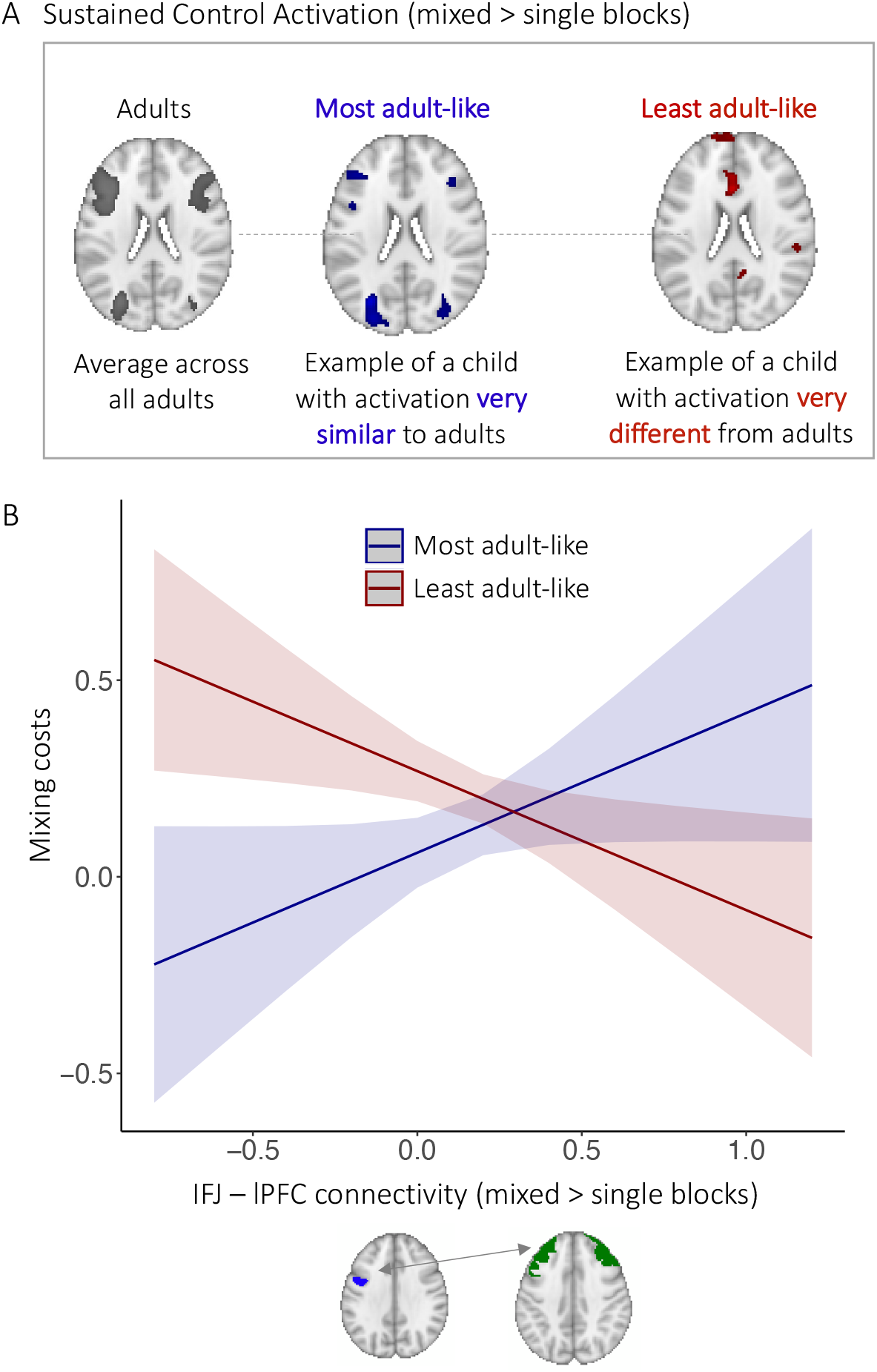
Interaction effect of deviation score and connectivity on accuracy mixing costs in children. (A) Sustained control activation (mixed > single) across adults (p < .05, FWE-corrected) used as reference for estimating how similar an individual child’s activation pattern was to this adult pattern. An example of a child showing more adult-like activation for the contrast shown in blue (p < .001, uncorrected). An example of a child showing less adultlike activation for the contrast shown in red (p < .001, uncorrected). (B) Interaction effect. Children who showed less adult-like activation patterns (red line) showed a negative association between connectivity increases from single to mixed block and mixing costs, such that greater increases in connectivity were associated with lower costs (i.e., better performance). Children showing more adult-like activation patterns (blue line) showed a positive association between connectivity increases and mixing costs, such that greater increases in connectivity were associated with greater mixing costs (i.e., worse performance).

In contrast, for children who showed less adult-like sustained control activation (i.e., more positive deviation scores), accuracy mixing costs and connectivity increases were *negatively* correlated. Thus, for children who deviated more from the expected sustained activation pattern, stronger incorporation of the lPFC into the control network—as evidenced by heightened connectivity with IFJ in response to an increase in the need for sustained control—was related to better performance. Similar results were obtained regarding IFJ–mPFC connectivity, albeit with the interaction of deviation scores and connectivity increases at a trend level (β = −0.15, p = .1). Neither the connectivity between the IFJ and the left AG nor the right AG showed such a relationship with performance.

Taken together, these results suggest that greater lPFC involvement represents an adaptive alternative to implement task-switching affordances among children whose FPN activation patterns deviate more from the adult pattern.

## 4. Discussion

In the present study, we sought to shed light on the development of sustained and transient control processes by taking a closer look at (a) age group differences between children aged 8 to 11 years and adults in performance, brain activation and connectivity related to transient and sustained task-switching demands; and (b) individual differences in the association between task-switching proficiency, neural activation and connectivity among children. Our results indicated that children showed greater mixing and switch costs than adults. While children engaged the same frontal and parietal regions during task switching, they upregulated brain activation in these areas to a lesser extent than adults. Age differences were more pronounced for mixing costs and sustained control activation, in line with the notion that brain areas and circuits associated with sustained control mature more slowly than those associated with transient control (Bunge and Wright, 2007; Cepeda et al., 2001; Crone et al., 2006a, 2004; Reimers and Maylor, 2005). Task-based connectivity analyses revealed that connectivity between the IFJ and the lPFC increased with greater sustained control demands in children. Critically, the association between task-related connectivity and task-switching performance was linked to how dissimilar the activation pattern of an individual child was to the typical adult pattern: Children who showed less adult-like activation in response to sustained control demands showed better performance with greater IFJ–lPFC connectivity increases, whereas children with a more adult-like activation pattern showed the opposite association, that is, lower performance with greater IFJ–lPFC connectivity increases. This finding suggests that children whose FPN is less mature make adaptive use of an alternative neural implementation of cognitive control to cope with increases in sustained switching demands.

### 4.1 Age differences in task-related activation and modulation by switching demand

Age differences were more pronounced in sustained control processes than in transient control processes, at both behavioral and neural levels. These results suggest that sustained control processes such as task-set selection (Chevalier et al., 2018; Chevalier and Blaye, 2009; Emerson and Miyake, 2003), maintenance, and monitoring (Braver et al., 2003; Pettigrew and Martin, 2016; Rubin and Meiran, 2005) follow an extended maturational path, potentially continuing into adolescence (Cepeda et al., 2001; Crone et al., 2006a, 2004; Huizinga and van der Molen, 2007; Reimers and Maylor, 2005). Of note, successful task selection depends on the ability to processes relevant task cues in the environment. A recent study using eye tracking during task switching revealed that adults and older children (8.5–12 years) first focused on the cue indicating the currently relevant task, whereas younger children (3–8 years) were more likely to first focus on the target (Chevalier et al., 2018). While these findings suggest that the children in the present study were old enough to process and select the goal-relevant task cue, the targets (face, scene, object) in our paradigm were presented at a different location on every trial. This trial-by-trial spatial reconfiguration requiring that participants search for the relevant stimulus on every trial may have drawn focus away from successful cue identification and undermined the successful engagement of sustained control.

At the same time, age differences in transient control were less pronounced than those in sustained control (behaviorally and in ROI analyses), or even absent in some analyses (whole-brain and meta-analysis ROIs). It is worth noting that previous findings regarding age differences in behavioral switch costs are inconsistent, with some studies reporting comparable switch costs in children and adults (Anderson et al., 2001; Luca et al., 2003; Reimers and Maylor, 2005) and others showing higher switch costs in children (Crone et al., 2006a; Huizinga et al., 2006). These discrepancies may be related to different facets of task switching, such as the ability to apply hierarchical rules beyond increased demands on working memory capacity (Unger et al., 2016) or the number of response options, even under low working memory demands (Bauer et al., 2017). While each being crucial to successful task switching, these facets may be targeted to different degrees across task-switching paradigms, thereby showing diverging developmental trajectories and different patterns of age differences in switch costs.

One potential explanation for the dissociation of age differences patterns between sustained and transient control is the concomitant difference in the developmental pace of brain regions activated for sustained or transient control demands. In our task, several frontal and parietal regions (i.e., the IFJ, SPL, and dACC) were recruited for both sustained and transient control; however, the dlPFC was exclusively modulated by sustained control demands. Previous research has shown that the dlPFC is among the latest brain areas to mature during childhood and adolescence (Sowell et al., 1999; Sydnor et al., 2021; but see Fuhrmann et al., 2022). Critically, an entire network’s efficiency can be hindered by a specific node, such that connectivity within a network is lower if a node of the network is lesioned, while networks that do not include this lesioned node are not affected in their efficiency (Nomura et al., 2010). Thus, while the regions associated with both transient and sustained control processes may be relatively more functional during transient control, they might be more strongly limited by the insufficient activation in the co-recruited dlPFC region during sustained control.

### 4.2 Age differences in task-based connectivity during task switching

Based on previous research demonstrating that connections within and between control networks continue to develop into adolescence (Fair et al., 2009, 2007; Luna et al., 2015; Marek et al., 2015), we expected children to show smaller increases in connectivity with greater sustained control demands, reflecting the ongoing development of these task-related networks. To the contrary, we found that children showed increased connectivity of the IFJ to regions that did not show reliable activation increases across children and adults in our task. One of these regions was a large PFC cluster with a lateral portion of the PFC, including the frontal pole/aPFC and middle frontal gyrus, and a medial portion, including the superior frontal gyrus and paracingulate gyrus. These results are consistent with a previous developmental connectivity study that also showed greater integration of the aPFC in a frontoparietal control network during task switching in children compared to adults (Ezekiel et al., 2013).

The lPFC has been suggested to follow a hierarchical organization, such that more posterior lPFC regions are associated with sensory-motor control, dorsolateral regions with contextual control, and the anterior regions with temporal control (Badre, 2008; Badre and D’Esposito, 2007; Badre and Nee, 2018; Nee and D’Esposito, 2016). The present results showing increased connectivity between the IFJ and a cluster stretching from posterior to anterior lPFC raises the question whether control hierarchies in the lPFC are less clearly established in children than in adults (Bunge and Zelazo, 2006; Unger et al., 2016), such that children might recruit higher-order control regions for lower-order demands (cf. Crone and Steinbeis, 2017). This idea is consistent with the protracted behavioral differentiation of cognitive control process in development (e.g., Akshoomoff et al., 2018; Best and Miller, 2010; Brydges et al., 2014; Shing et al., 2010) and the less flexible recruitment of control strategies based on current control demands in younger children (e.g., Chevalier et al., 2019; Chevalier, 2015).

The maturity of children’s pattern of activation in response to sustained control demands moderated the relation between IFJ–lPFC connectivity and performance. More precisely, in children showing less adult-like activation, increased IFJ–lPFC connectivity had a positive effect on performance, suggesting that upregulating lPFC involvement with increased sustained control demands functioned as a performance-enhancing alternate neural strategy in this group of children.

lPFC activation and connectivity supports the management (Lara and Wallis, 2015; Miller et al., 2018; Sala and Courtney, 2007) and selection (Badre, 2012; Chatham and Badre, 2015; D’Ardenne et al., 2012) of multiple task-sets in adults. In this context, our findings indicate that additional recruitment of a region supporting task selection and management is more likely to lead to reduced mixing costs in children, for whom core task switching regions are not yet sufficiently mature to be flexibly activated. This idea is in line with recent discussions of meta-control, broadly defined as control processes that monitor and regulate other control processes (Eppinger et al., 2021) and determine when, how much, and what type of control to exert (Lieder et al., 2018). Children’s meta-control improves considerably in middle childhood (Chevalier, 2015; Chevalier and Blaye, 2016; Niebaum et al., 2021; Schuch and Konrad, 2017). The lPFC has been suggested to play a key role in meta-control (Eppinger et al., 2021; Ruel et al., 2021), suggesting that children who showed less mature patterns of task-specific neural activation may have achieved successful task-switching performance by increased reliance on meta-control monitoring and/or regulation.

Alternatively or additionally, children struggle particularly with the management of multiple task-set representations (Crone et al., 2006a, 2004), a key component of sustained control during task switching (Braver et al., 2003; Pettigrew and Martin, 2016; Rubin and Meiran, 2005). Especially when task-set representations are less distinct or unavailable, as might be expected of children showing less adult-like activation patterns, they might benefit from additional resources to accomplish this management process. The lPFC is ideally suited to provide this additional support due to its role in managing attention between internal information (e.g., maintained task-set representations) and external (stimulus) information (Burgess et al., 2007).

In contrast, in children showing more adult-like sustained activation, increased IFJ–lPFC connectivity in mixed blocks was negatively associated with task performance. Interestingly, adults showed a similar brain-behavior relationship (albeit not statistically significant), such that greater increases in connectivity were associated with greater mixing costs. We surmise that once the core regions involved in task switching are relatively more mature, involving additional peripheral regions may make their operation less efficient. Indeed, under certain circumstances, increased control and lPFC involvement can hinder learning and task performance, in particular during stimulus-driven responses and sequence learning (Galea et al., 2010; Kruschke, 2003; Thompson-Schill et al., 2009). Increasingly specific and efficient networks (Chevalier et al., 2019; Dreher and Berman, 2002; Fair et al., 2009, 2007; Zhang et al., 2021) and more flexible adjustments of strategies (Chevalier et al., 2015) can give rise to more adult-like performance. However, as our results suggest, they also harbor room for selecting inappropriate or inefficient strategies due to ongoing finetuning of brain networks and the neurocognitive processes they support. Critically, developmental changes in brain activation and connectivity might show non-linear and possibly non-monotonic relations to behavior that differ between and vary within individuals (Lautrey, 2003; Li and Lindenberger, 2002; Wendelken et al., 2017), further contributing to the complexity of developmental transitions in the neural machinery supporting task switching behavior. Longitudinal studies are needed to understand how brain structure, activation, and connectivity develop interactively with behavior to enable efficient task switching in the course of development. Such studies need to investigate how neural activation at one timepoint is related to connectivity at another, and importantly, how these longitudinal lead-lag relations, and individual differences therein, correlate with, and possibly predict changes in behavior.

### 4.3 Conclusion

Taken together, the presented data suggest that sustained and transient control processes during task switching follow different developmental trajectories, such that transient processes approximate adult levels relatively earlier than sustained processes do. A potential mechanism giving rise to children’s task-switching difficulties might be inefficient upregulation of activation in response to increased control demands, especially for sustained control. However, increased connectivity between the IFJ and the lPFC with greater sustained control demands might offer an alternative, and possibly developmentally earlier mechanism to manage these demands, at least in some children. These findings point to a complex pattern of brain-behavior relationships during task switching in which children can cope with increased sustained control demands in at least two ways, either by upregulating activation in brain regions engaged by adults or by enhancing connectivity with brain areas involved in the maintenance and management of multiple task sets. The longitudinal study of these patterns will reveal how these strategies shift in the course of development and how they are influenced by experience.

## Acknowledgements

We acknowledge financial support by the Max Planck Institute for Human Development and by the DFG Priority Program SPP 1772 “Human performance under multiple cognitive task requirements: From basic mechanisms to optimized task scheduling” (Grant No. FA 1196/2-1 to Y.F.). S.A.S. and N.K. are doctoral fellows of the International Max Planck Research School on the Life Course (LIFE; http://www.imprs-life.mpg.de). The authors thank Julia Delius for editorial assistance and helpful comments. Communication regarding this article should be directed to S.A.S, or Y.F..

