## SupplementaryMaterial for "Does prefrontal connectivity during task switching help or hinder children’s performance?"

### Supplementary Material

| | Hemisphere | Region | Mixed > Single Peak | Switch > Repeat Peak | Size (Voxels) |
|---|---|---|---|---|---|
| Overlap | left | IFJ | -40 2 32 | -40 0 40 | 240 |
| | | SPL | -28 -58 46 | -28 -50 44 | 303 |
| | | dACC | -4 14 52 | -8 8 52 | 173 |
| Sustained | left | dlPFC | -48 22 32 | | 952 |
| | right | dlPFC | 50 32 24 | | 534 |
| | | IFJ | 42 4 30 | | 352 |
| | | SPL | 30 -56 48 | | 299 |
| Transient | left | Precuneus | | -16 -68 24 | 489 |
| | right | Precuneus | | 22 -58 26 | 390 |

*Supplementary Table 1*: Regions of interest (ROIs) showing activation under sustained (mixed > single blocks) and transient (switch > repeat trials) control demand (Overlap), exclusively under sustained control demand and exclusively under transient control demand, respectively. Coordinates (X Y Z) in Montreal Neurological Institute space. Size in voxels (2 mm x 2 mm x 2 mm). Abbreviations: IFJ: inferior frontal junction, SPL: superior parietal lobe, dACC: dorsal anterior cingulate cortex, dlPFC: dorsolateral prefrontal cortex.

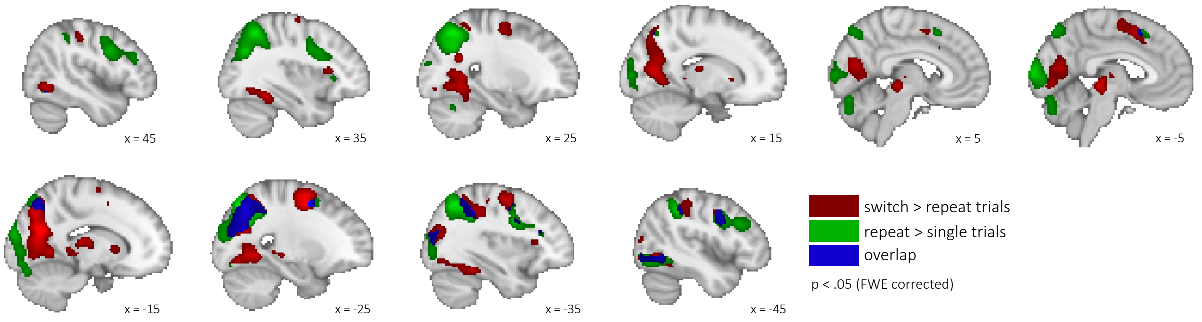


*Supplementary Figure 1: Univariate activation across age groups in control GLM.* Switch > repeat trials depicted in red, repeat > single trials in green, and overlap in blue. N = 142 (53 adults, 89 children), $p < .05$ FWE-corrected, $k = 50$.

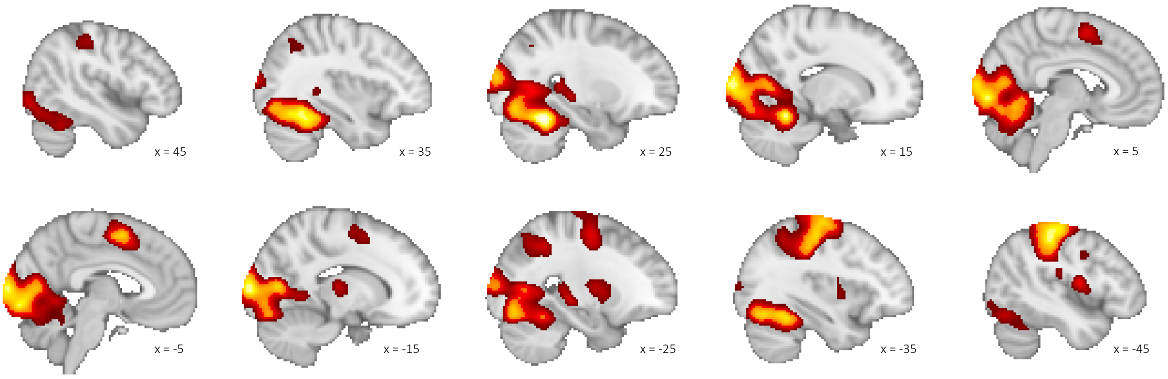


*Supplementary Figure 2a: Univariate activation in single blocks.* Activation in single blocks (vs. baseline), N = 142 (53 adults, 89 children), $p < .05$ FWE-corrected, k = 50.

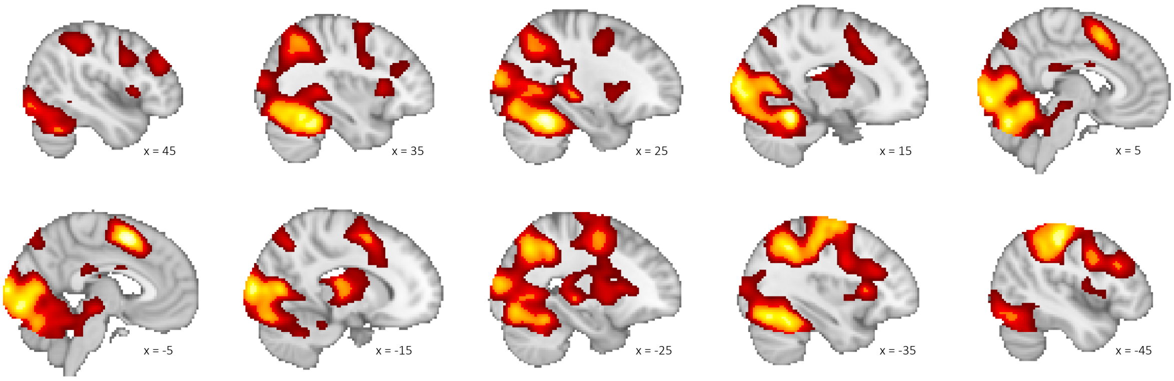


*Supplementary Figure 2b: Univariate activation in mixed blocks.* Activation in mixed blocks (vs. baseline), N = 142 (53 adults, 89 children), $p < .05$ FWE-corrected, k = 50.

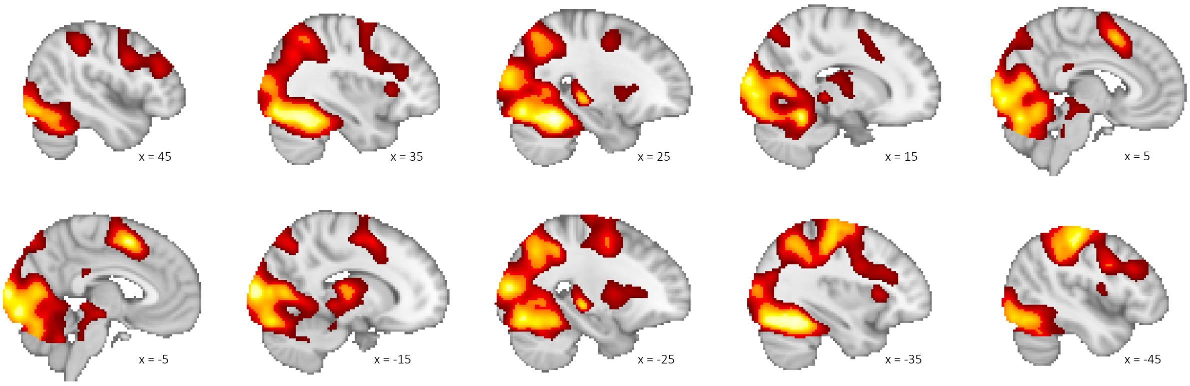


*Supplementary Figure 2c: Univariate activation on repeat trials.* Activation on repeat trials (vs. baseline), N = 142 (53 adults, 89 children), $p < .05$ FWE-corrected, k = 50.

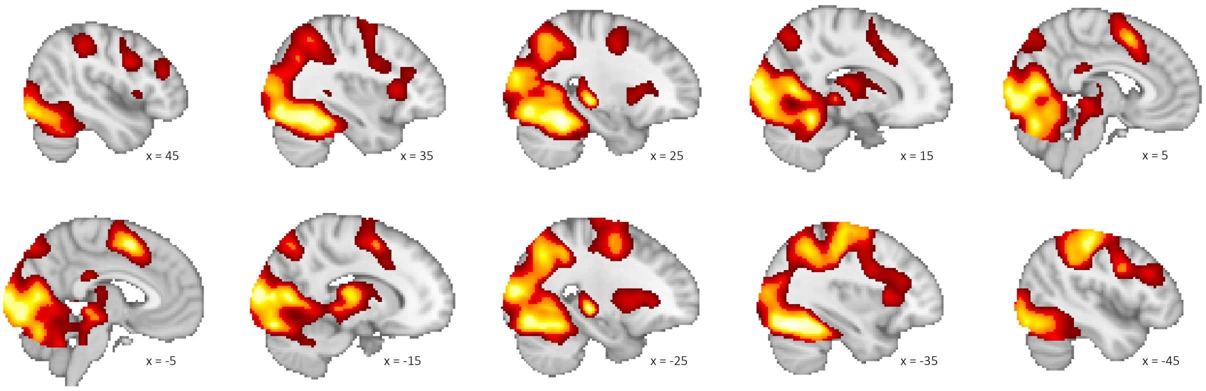


*Supplementary Figure 2d: Univariate activation on switch trials.* Activation on switch trials (vs. baseline), N = 142 (53 adults, 89 children), p < .05 FWE-corrected, k = 50.

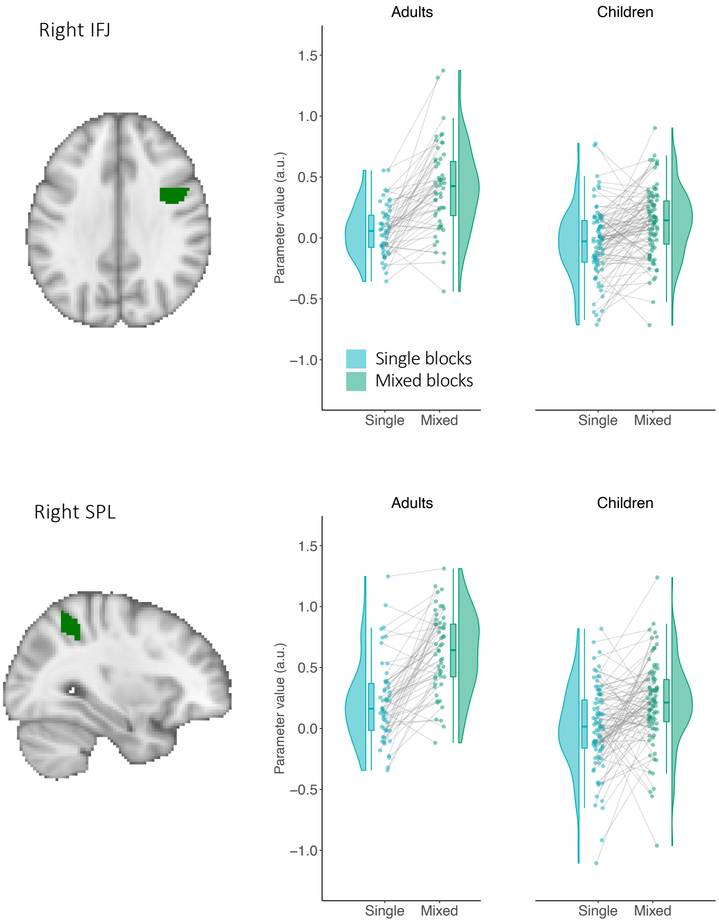


Supplementary Figure 3: *Age differences in activation due to sustained control demand by ROI.* Extracted parameter estimates for single blocks (blue) and for mixed blocks (green). Children showed less upregulation of sustained activation in the right IFJ and SPL.

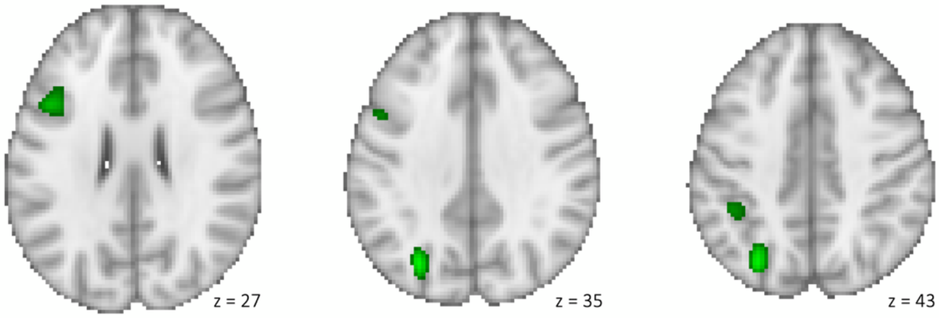


*Supplementary Figure 4a: Whole brain age differences*. Clusters showing greater sustained control activation (mixed > single contrast) in adults than in children ($p < .05$ FWE-corrected, k=50).

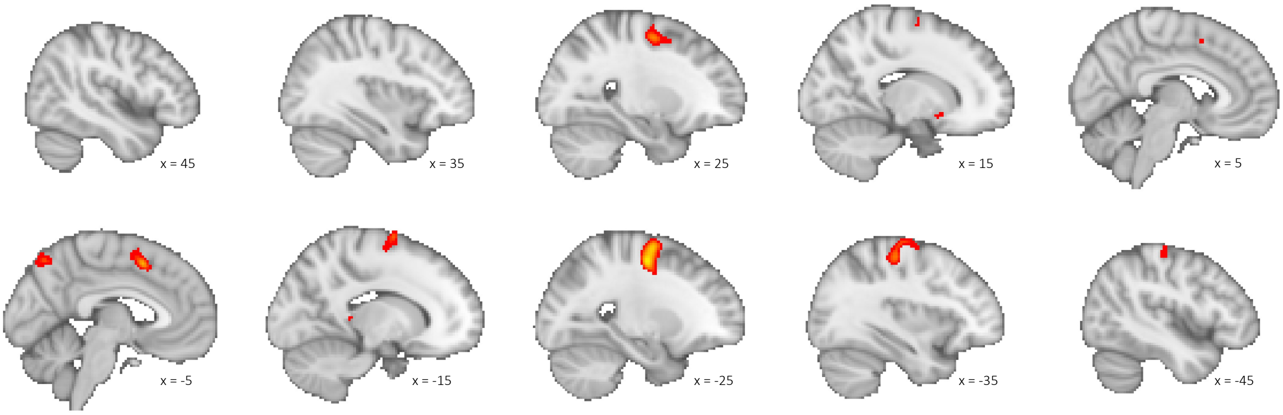


*Supplementary Figure 4b: Whole brain age differences*. Clusters showing greater transient control activation (switch > repeat trials) in adults than in children ($p < .001$ uncorrected, k=50). No clusters survived correction for multiple comparisons with $p < .05$ FWE-corrected.

| **whole sample (adults & adolescents)** | | MNI coordinates | | | Age comparison analysis (t-values) | |
|---|---|---|---|---|---|---|
| Hemisphere | Brain region | X | Y | Z | sustained activation | transient activation |
| left | Inferior parietal lobule/superior parietal lobule/precuneus | -40 | -42 | 44 | -5.03* | -1.77 |
| right | Inferior parietal lobule/superior parietal lobule/precuneus | 40 | -42 | 44 | -3.04* | -0.63 |
| L | Precentral gyrus/middle frontal gyrus | -44 | 6 | 32 | -4.74* | -2.08 |
| L | Pre-supplementary motor area/superior frontal gyrus | -4 | 20 | 48 | -1.73 | -1.64 |
| R | Precentral gyrus/middle frontal gyrus | 44 | 8 | 30 | -3.31* | -0.099 |
| L | Middle frontal gyrus/precentral gyrus | -26 | -6 | 54 | -3.43* | -3.96* |
| L | Thalamus/putamen/caudate | -10 | -16 | 10 | -1.53 | -1.49 |
| R | Middle frontal gyrus | 30 | 0 | 58 | -1.95 | -2.93* |
| R | Insula | 34 | 24 | -2 | -2.55* | -1.52 |
| L | Insula | -32 | 22 | 2 | -3.27* | -0.91 |

*Supplementary Table 2*: Age comparison analysis results of spherical ROIs (6mm radius) based on regions showing switching related activation across studies with adults and adolescents in Zhang et al., 2021. T-tests were conducted separately for the difference between adults and children in sustained activation (i.e., difference in activation between single and mixed blocks) and transient activation (i.e., difference in activation between repeat and switch trials). Higher t-values represent a greater difference between condition (i.e., greater upregulation of activation) in adults. Note: * denote FDR-corrected $p < .05$

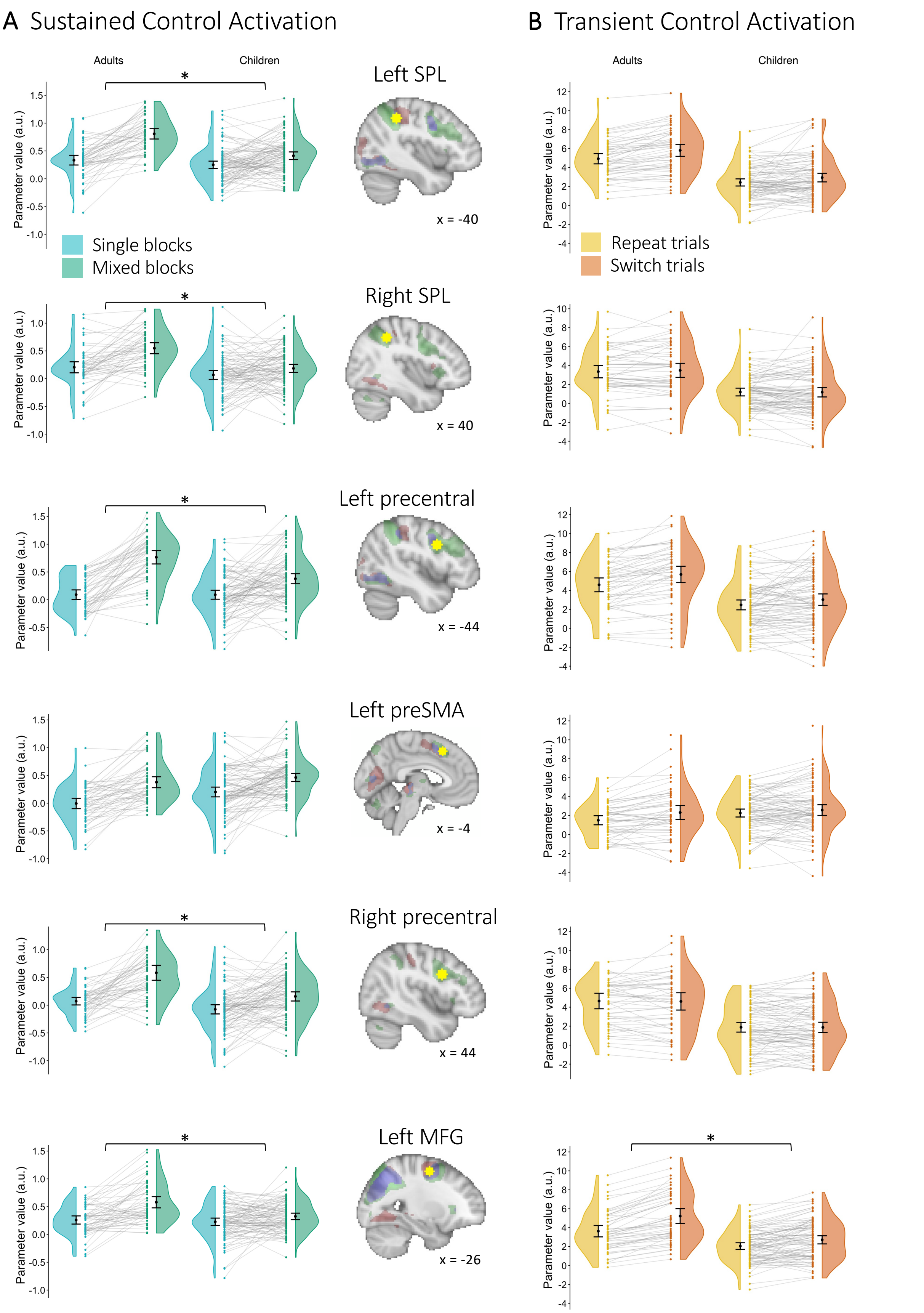


*Supplementary Figure 5: Age differences in activation due to sustained and transient control demands in meta-analysis ROIs (Zhang et al. 2021).* Figure description on next page.

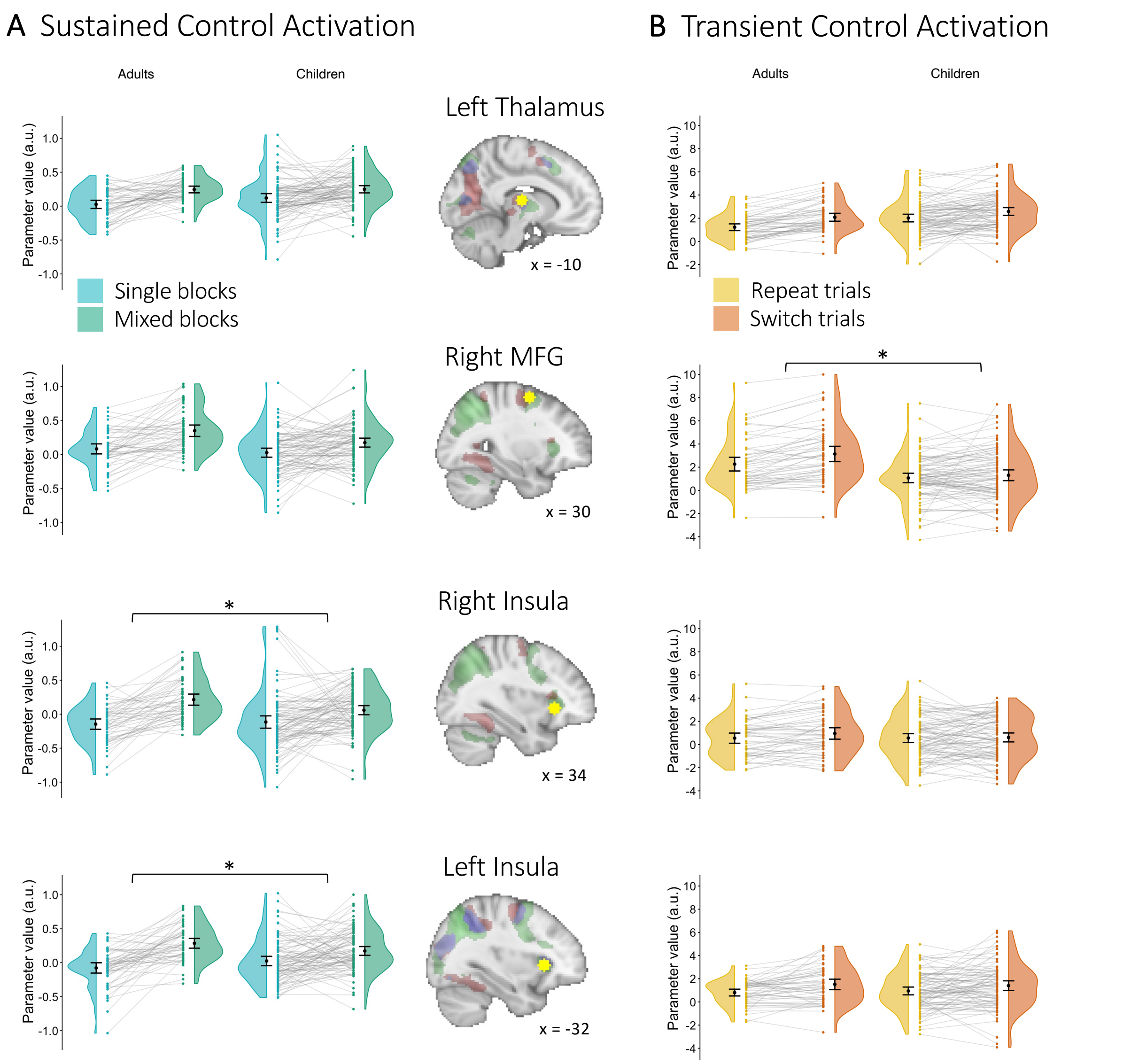


*Supplementary Figure 5 (continued): Age differences in activation due to sustained and transient control demands in meta-analysis ROIs (Zhang et al. 2021). ).* Extracted parameter estimates from ROIs 6mm spheres around the peaks found by the meta-analysis across adults and adolescents for (A) single blocks (blue) and for mixed blocks (green) and (B) for repeat trials (yellow) and for switch trials (orange). Spheres are displayed in yellow overlain on the sustained/transient/ overlap activation across adults and children in the present study in green, red, and blue, respectively. * denote FDR-corrected $p < .05$

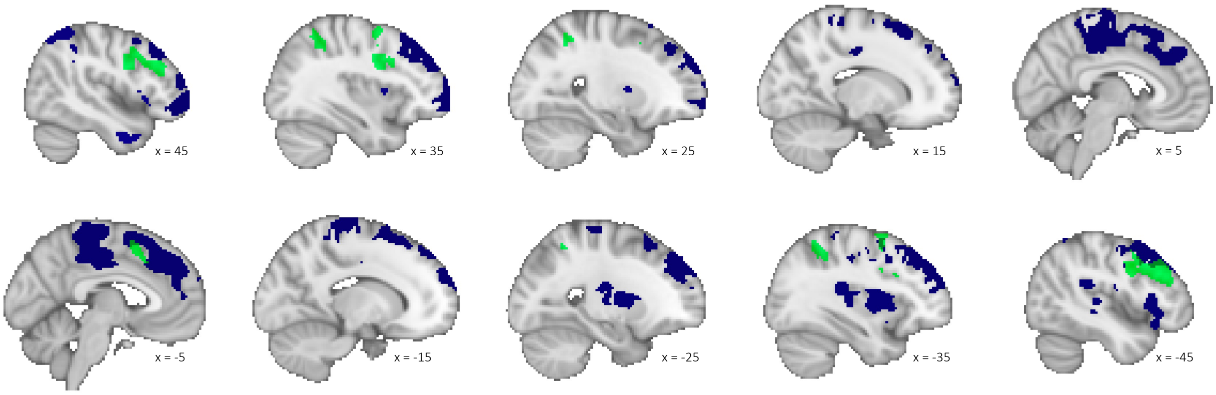


*Supplementary Figure 6:* Sustained and Overlap ROIs (green) and clusters showing greater connectivity difference (mixed > single blocks) in children than adults (blue). The connectivity clusters do not substantially overlap with the ROIs based on task-related activation.

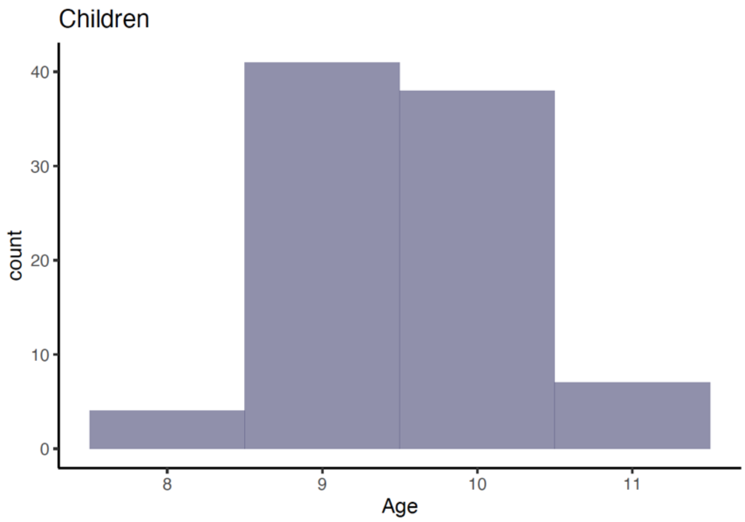


*Supplementary Figure 7*: Distribution of age across the age span of children included in the present manuscript.
